# Enzymatic assay for UDP-GlcNAc and its application in the parallel assessment of substrate availability and protein O-GlcNAcylation

**DOI:** 10.1101/2023.03.18.533286

**Authors:** Marc Sunden, Divya Upadhyay, Rishi Banerjee, Nina Sipari, Vineta Fellman, Jukka Kallijärvi, Janne Purhonen

**Affiliations:** Folkhälsan Research Center, Helsinki, Finland; Stem Cells and Metabolism Research Program, Faculty of Medicine, University of Helsinki, Helsinki, Finland; Viikki Metabolomics Unit, University of Helsinki; Finland; Children’s Hospital, Helsinki University Hospital, Finland, and Pediatrics, Department of Clinical Sciences Lund, Lund University

## Abstract

O-linked N-acetylglucosaminylation (O-GlcNAcylation) is a ubiquitous and dynamic yet still relatively poorly understood non-canonical glycosylation of intracellular proteins. Several vital branches of metabolism converge at the hexosamine biosynthetic pathway (HBP) to produce the substrate for protein O-GlcNAcylation the uridine diphosphate N-acetylglucosamine (UDP-GlcNAc). Availability of this metabolite is considered a key regulator of O-GlcNAcylation. Yet UDP-GlcNAc concentrations are rarely reported in studies exploring the HBP and O-GlcNAcylation, most likely because the methods to measure it restrict to specialized chromatographic procedures. To overcome this limitation, we introduce here an enzymatic method to quantify cellular and tissue UDP-GlcNAc. The method is based on O-GlcNAcylation of a substrate peptide by recombinant O-linked N-acetylglucosamine transferase (OGT) and detection of the modification with a specific antibody. The assay can be performed in dot blot or microplate formats. The key to successful assay was the removal of strong inhibition of OGT by the reaction side product, uridine diphosphate (UDP). We applied the assay to provide the first systematic report of UDP-GlcNAc concentrations in mouse tissues and cultured cells. Furthermore, we show how changes in UDP-GlcNAc levels correlate with O-GlcNAcylation and the expression of OGT and O-GlcNAcase (OGA).

## Introduction

The hexosamine biosynthetic pathway (HBP) consumes uridine triphosphate (UTP), glucose, glutamine, and acetyl-CoA to produce two N-acetylated amino sugars coupled to uridine diphosphate (UDP): UDP-N-acetylglucosamine (UDP-GlcNAc) and UDP-N-acetylgalactosamine (UDP-GalNAc).^1^ In addition to being a biosynthetic precursor for glycan chains of extracellular proteins, UDP-GlcNAc is the substate for the monomeric O-linked N-acetylglucosaminylation (O-GlcNAcylation) of serine and threonine residues of intracellular proteins. The sole enzyme performing this specific glycosylation in mammalian cells is the O-linked N-acetylglucosamine transferase (OGT). Likewise, only one enzyme, the O-GlcNAcase (OGA), removes this modification. While seemingly simple two-enzyme regulation, an increasing number of studies showing altered protein O-GlcNAcylation in various pathological conditions and experimental models implicate O-GlcNAc as a highly dynamic protein post-translational modification.^1^

Many reviews have postulated UDP-GlcNAc as the central regulatory metabolite governing protein O-GlcNAcylation.^2–5^ Indeed, the HBP integrates nucleotide, carbohydrate, amino acid and fatty acid metabolism, making UDP-GlcNAc a sensible regulatory node for energy metabolism. Nevertheless, cellular or tissue concentrations of UDP-GlcNAc have rarely been reported even in many seminal studies on O-GlcNAcylation. Based on limited literature, tissue UDP-GlcNAc concentrations range from 10 to 35 μM in the skeletal muscle to ~150 μM in the liver,^6–10^ while OGT requires only 0.5-5 μM UDP-GlcNAc for its half-maximal activity,^11–13^ contradicting the UDP-GlcNAc availability as a sensitive regulator of OGT activity at least in some tissues. However, UDP-GlcNAc does not equally distribute within a cell but is actively concentrated into the endoplasmic reticulum and Golgi apparatus.^14^ Unfortunately, little is known about the effective UDP-GlcNAc concentration in different cellular compartments.

Traditional methods to measure nucleotide sugars involve chromatographic procedures such as liquid chromatography or capillary electrophoresis.^10,15,16^ These methods require expensive special equipment and expertise, and thus are not easily applicable in a general research laboratory. Moreover, the near-identical chromatographic elution profile, reactivity, physical properties, and the identical molecular mass of the nucleotide sugar epimers, make their separate quantification (e.g. UDP-GlcNAc from UDP-GalNAc) challenging using these methods.^17^ For these reasons, UDP-GlcNAc and UDP-GalNAc concentrations are typically reported as a sum of these two metabolites (UDP-N-acetylhexosamines, UDP-HexNAc). The biological functions of the two UDP-HexNAcs are, however, distinct.

To date, only one enzymatic assay for UDP-GlcNAc has been published.^18^ This assay is based on NAD^+^-dependent UDP-GlcNAc dehydrogenase from *Methanococcus maripaludis*. As the assay relies on direct monitoring of NADH generated, it is intrinsically limited in sensitivity. The authors showed, however, sufficient sensitivity to measure UDP-GlcNAc from yeast and Hela cells. Nevertheless, to our knowledge, no further use of this method has been reported during the 14 years since its publication.

Here, we took advantage of the high affinity of OGT for UDP-GlcNAc to develop an enzymatic assay for UDP-GlcNAc. We utilized the assay to measure UDP-GlcNAc concentrations in different mouse tissues and cultured cells. Finally, we examined the relationships between UDP-GlcNAc levels, protein O-GlcNAcylation, and the expression of OGT and OGA.

## Results

### Development of enzymatic assay for UDP-GlcNAc

After unsuccessful attempts to quantify UDP-GlcNAc with the previously reported enzymatic assay^18^ (Supplementary Fig. 1) and competitive enzyme-linked lectin-binding approach (Supplementary Materials and Methods), we looked into methods to measure glycosyltransferase activity of OGT,^19,20^ and whether these methods could be converted for the measurement of UDP-GlcNAc. Given the reported high affinity of OGT for UDP-GlcNAc and multiple options to detect O-GlcNAcylated proteins, we reasoned that OGT-mediated O-GlcNAcylation could, in theory, be utilized to develop an assay for UDP-GlcNAc. For proof-of-principle experiments, we set up a simple assay scheme involving human recombinant OGT fragment, a GlcNAc-acceptor peptide crosslinked to bovine serum albumin (BSA), limiting concentrations of UDP-GlcNAc, and dot blotting onto PVDF membrane to capture the peptide-BSA complex for immunodetection of O-GlcNAcylated residues (Figure 1A). The acceptor peptide derives from human casein kinase 2 and is one of the most efficient substrates for O-GlcNAcylation reported.^12^ We reasoned that for the assay to work, we would need to remove interference by UDP, especially from biological samples. UDP is a reaction product in O-GlcNAcylation and a potent inhibitor (IC_50_ < 1 μM) of OGT.^11,21^ To do this, we included alkaline phosphatase in the assay reactions. The pyrophosphate group in free nucleotides is exposed to hydrolysis by phosphatases, whereas in UDP-GlcNAc it is not as it is protected by the glucosamine and ribose moieties. For immunodetection of O-GlcNAc, we chose the mouse monoclonal antibody RL2.^22^ While several other GlcNAc-specific antibodies and lectins exist, many of them cross-react with glycosylated proteins in blocking reagents such as skimmed milk and BSA or tend to give high background for other reasons. This is our experience and also reported in the literature.^19,23^

**Figure 1.**
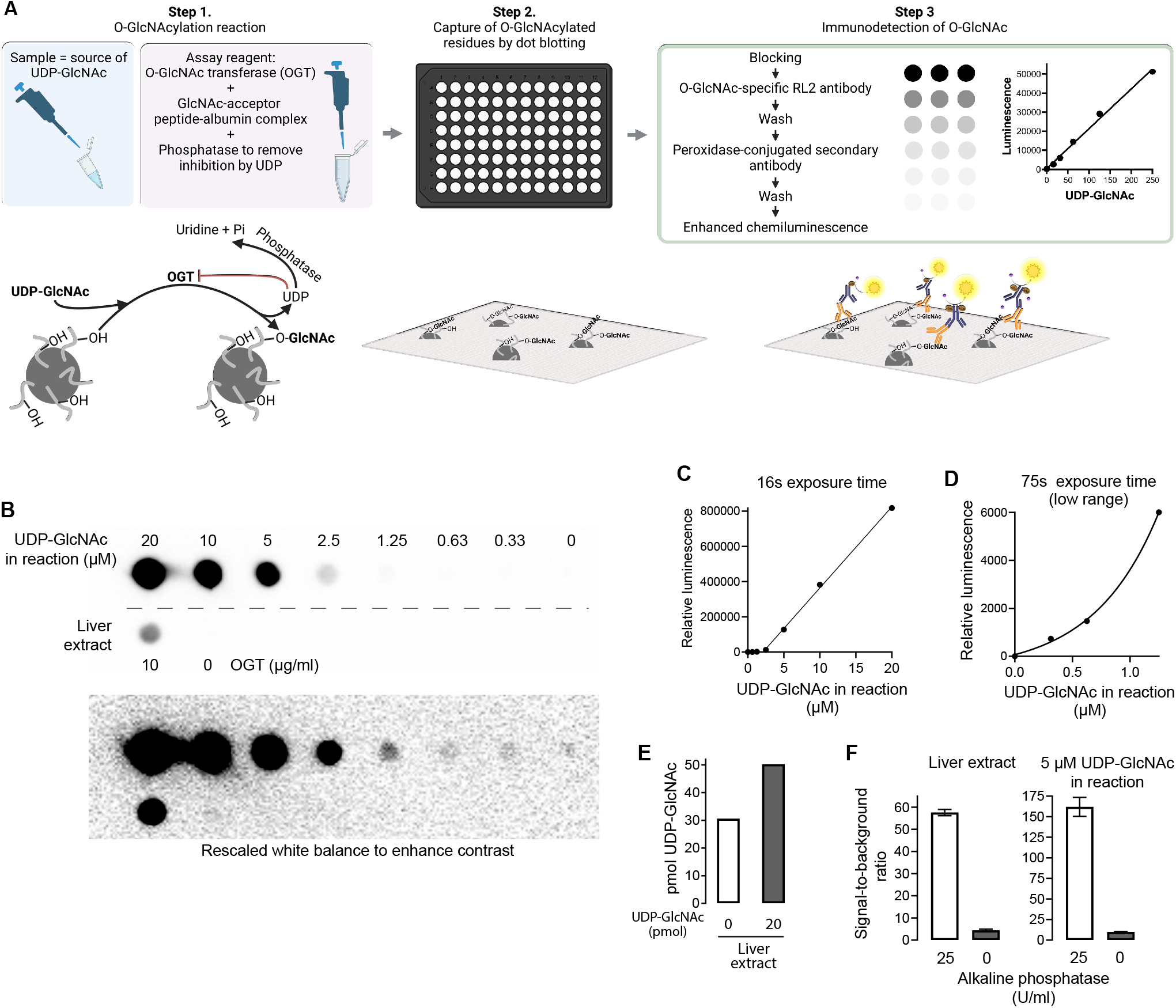
Enzymatic assay for UDP-GlcNAc with dot blot detection. (**A**) A schematic presentation of the assay principle. (**B**) A representative detection of O-GlcNAcylated peptides by dot blotting, RL2 monoclonal antibody, and chemiluminescence. (**C**) Standard curve showing UDP-GlcNAc concentration-dependent increase in signal. (**D**) Standard curve covering 0 to 1.25 μM UDP-GlcNAc after extended camera exposure time. (**E**) Measured UDP-GlcNAc amount from a liver extract with and without addition of 20 pmol exogenous analyte. (**F**) Effect of UDP degradation (by alkaline phosphatase) on measured signal from liver extract and purified UPD-GlcNAc standard sample. The error bars represent SEM of three technical replicates.

The initial assay concept was successful with purified UDP-GlcNAc standards and, importantly, with liver tissue extract (Figure 1B-E). In the absence of OGT, the liver extract did not give any measurable signal, verifying complete removal of endogenous O-GlcNAcylated proteins (Figure 1B). Spiking the liver extract with a known amount of UDP-GlcNAc led to a corresponding increase in signal (Figure 1E), excluding major sample-derived interference. In line with our hypothesis, the inclusion of alkaline phosphatase proved to be indispensable for the assay (Figure 1F).

### Enzymatic UDP-GlcNAc assay in microplate format

While we found the dot blot format simple, quantitative and relatively sensitive, our CCD-camera based imaging system limited the dynamic range. Moreover, this assay format requires a dot blotting apparatus for quantitative results, a relatively rare piece of equipment nowadays. Because of these limitations, we modified the assay into 384-well microplate format (Figure 2A). We coated the wells of high protein-binding microplates with the GlcNAc-acceptor peptide-BSA complex. After blocking unoccupied protein-binding sites, we performed the O-GlcNAcylation reactions in the wells and utilized a typical direct ELISA-like detection using RL2 antibody and peroxidase-conjugated secondary antibody. We chose to develop the signal using Amplex UltraRed as the peroxidase substrate, giving a highly sensitive chemifluorescent read out. Figure 2B shows a standard curve generated from the end-point fluorescence values.

**Figure 2.**
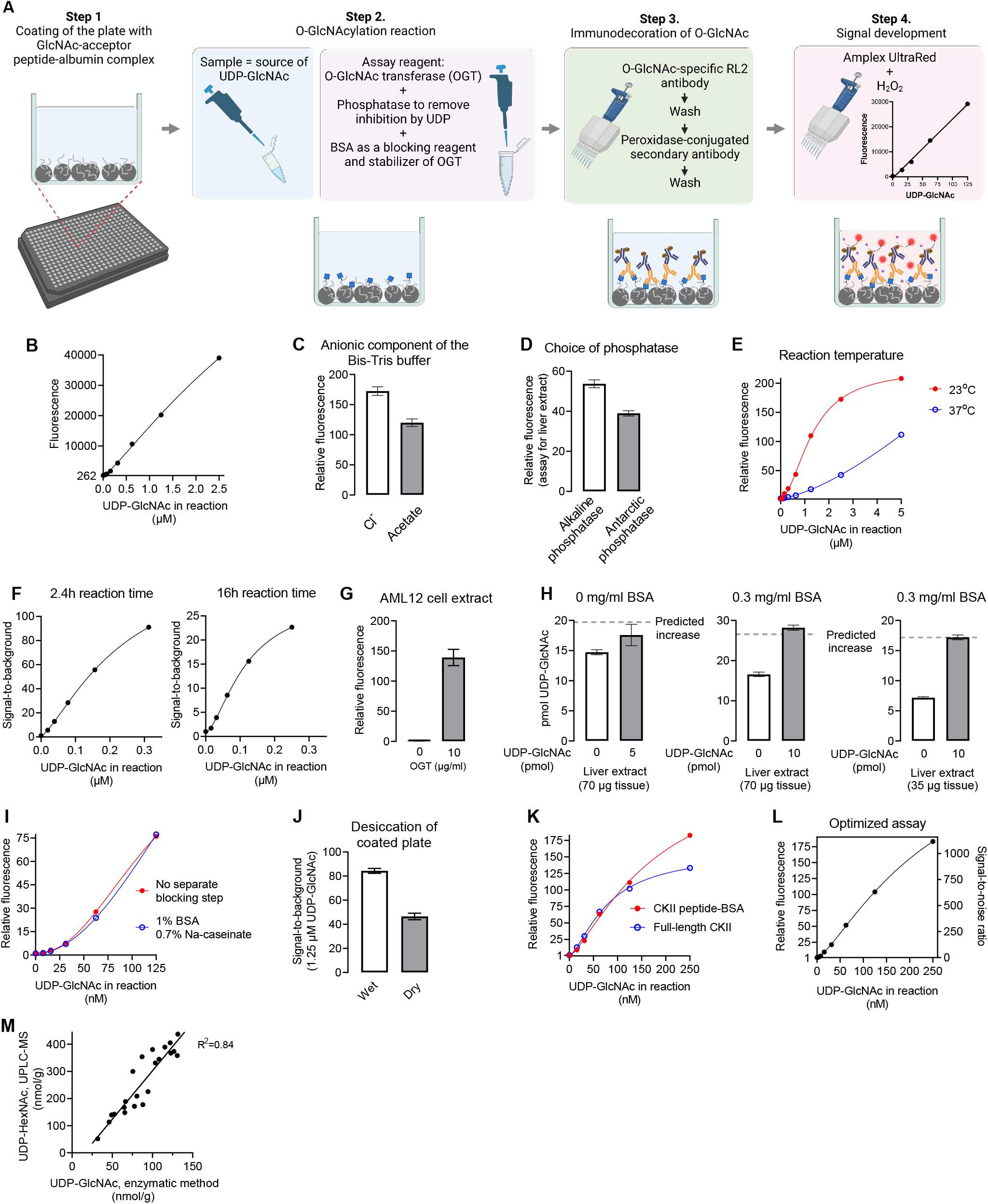
Enzymatic UDP-GlcNAc assay in microplate format. (**A**) A schematic presentation of the microplate assay for UDP-GlcNAc. (**B**) Standard curve generated from end-point fluorescence values. (**C**) Comparison of chloride and acetate as counter anions in Bis-Tris-based assay buffer. (**D**) Comparison of two phosphatases with different pH optimum on signal from liver extract. The assay buffer pH was 7.0. (**E**) Effect of O-GlcNAcylation reaction temperature on the assay performance. (**F**) Effect of reaction time on the sensitivity of the assay. (**G**) Verification of the specificity with a biological sample by omission of OGT from the assay reactions. (**H**) Effect of BSA in assay reactions on sample-related inhibition as assessed by spiking the samples with a known amount of UDP-GlcNAc. OGT concentration in these measurements was 10 μg/ml. (**I**) Dispensability of the separate blocking step. (**J**) Effect of desiccation of the coated plate. (**K**) Comparison of human casein kinase II and the peptide-BSA complex as GlcNAc acceptors in the assay reactions. (**L**) A representative standard curve of the optimized assay. (**M**) Correlation between UDP-GlcNAc (microplate assay) and UDP-HexNAc levels (UPLC-MS) in mouse liver samples. Bar graphs present mean and SEM of technical triplicates.

### Optimization of the assay

Using the dot blot and microplate formats of the assay, we set out to optimize the assay parameters for highest performance. The reported pH optimum for OGT activity lies between 6 to 7.5.^11^ Different buffers (Tris-Cl, Bis-Tris-Cl, Bis-Tris-acetate, HEPES-K, pH 7-7.5) did not have large impact on OGT activity under limiting substrate concentration (Supplementary Figure 2A-C and Figure 2C). OGT is highly sensitive to inhibition by potassium and sodium chloride according to some reports.^11,21^ We tested a HEPES-based buffer with near-physiological concentration of potassium (~90 mM) but without unphysiological chloride content. This buffer was compatible with the assay but inferior to plain Bis-Tris buffer that lacked added salt (Supplementary Figure 2B). When using Bis-Tris buffer, acetate proved an inferior counter anion in comparison to chloride (Figure 2C). For further experiments, we settled for 50 mM Bis-Tris (Cl^-^) buffer with pH 7.0. OGT does not require divalent cations for its activity,^24^ and, indeed, omission of Mg^2+^ had only a minor effect when assessing purified UDP-GlcNAc samples (Supplementary Fig. 1A). We, however, decided to include 5 mM Mg^2+^ to maximize alkaline phosphatase activity, which might become limiting when assessing complex biological samples. The selected pH 7.0 is not optimal for the alkaline phosphatase activity. However, a phosphatase with pH optimum close to 7 (New England BioLabs, Antarctic phosphatase) performed worse than alkaline phosphatase when assessing liver extracts (Figure 2D).

For the aforementioned experiments, we used a reaction time of 2h at room temperature. OGT has been reported to be relatively stable *in vitro* at temperatures below 30°C but to undergo rapid inactivation at 37°C.^21^ In the dot blot format, increase of reaction temperature to 37°C had only a minor buffer-dependent effect on the signal (Supplementary Figure. 1A). In the microplate format, however, the increase of reaction temperature to 37°C notably compromised the assay sensitivity (Figure 2E). This assay format difference suggests that GlcNAc-acceptor peptides, which were in excess and in solution in the dot blot format, help to stabilize OGT. We found that the GlcNAcylation reactions could be left to proceed overnight (~16h) for convenience (Figure 2F). However, the extended reaction time increased both the specific signal and the background, therefore decreasing sensitivity (Figure 2F).

Next, we assessed cell and liver extracts with the microplate format. Cell extracts gave measurable signal only when the assay reactions included OGT (Figure 2G), verifying the specificity. Surprisingly, the microplate format turned out to be more sensitive to sample-related inhibition than the dot blot format (Figure 2H). In the dot blot format, the reactions contained BSA (as a carrier for the GlcNAc-acceptor peptide). Therefore, we suspected that BSA might mitigate the sample-related inhibition. Indeed, this was the case (Figure 2H).

Due to the relatively high protein concentration (BSA, OGT, and alkaline phosphatase) in O-GlcNAcylation reactions, we tested the dispensability of the separate blocking step. This step turned out to be unnecessary (Figure 2I), and we omitted it from the final protocol.

For initial experiments, we used sodium carbonate bicarbonate buffer pH 9.6 to coat the microplates, but PBS worked equally well as the coating buffer (Supplementary Figure 2D). The coated plates did not tolerate desiccation for storage without loss of the assay sensitivity (Figure 2J). As an alternative GlcNAc acceptor, we tested coating with commercial human casein kinase 2 (composed of both α- and β-subunits). This GlcNAc acceptor gave essentially identical assay behavior to the peptide-BSA complex when UDP-GlcNAc concentrations were below 125 nM (Figure 2K). At higher concentrations, the signal plateaued earlier than with the peptide-BSA complex, suggesting that the GlcNAcylation sites became limiting.

Finally, we tested the lowest limit of quantification of the optimized assay (Figure 2L). Our microplate format reached the lowest limit of quantification (LLOQ) of 110 fmol (5.5 nM UDP-GlcNAc in reaction). The estimated lowest limit of detection (LLOD) was 40 fmol.

### UDP-GlcNAc concentrations in mouse tissues

To test the applicability of our method, we measured UDP-GlcNAc from several mouse tissues. Different nucleotide sugars extraction procedures have been reported in the literature ranging from acid (e.g. perchloric acid and trichloroacetic acid) to solvent extractions. ^10,17,18^ We chose MeOH-H_2_O-CHCl_3_ extraction, which has been shown to give close to 100% recovery for nucleotide sugars,^25^ and which we found compatible with the enzymatic method and ultra-performance liquid chromatography - mass spectrometry (UPLC-MS). For this extraction method to be compatible with the enzymatic method, we removed the residual MeOH from the aqueous phase by repeated CHCl_3_ or diethyl ether washes, or alternatively by drying the extracts. The recovery of exogenous UDP-GlcNAc spiked to kidney biopsies immediately after the tissue disruption was over 90% (Supplementary Figure 3). As an assay validation, we measured UDP-GlcNAc from 23 mouse liver samples from which we had existing UDP-HexNAc data (determined by UPLC-MS). Assuming unrestricted cellular epimerization between UDP-GlcNAc and UDP-GalNAc, the levels of these two metabolites should show high correlation. This was indeed the case, and the UDP-GlcNAc concentrations strongly correlated with the total UDP-HexNAc levels (Figure 2M).

Table 1 shows UDP-GlcNAc concentrations in the mouse liver, kidney, heart, skeletal muscle, and brain. Of these tissues, the liver had the highest UDP-GlcNAc concentration (240 pmol/mg). This concentration was of similar magnitude to previously reported values (125-150 pmol/mg).^6^ The skeletal muscle contained the smallest amount of UDP-GlcNAc (14 pmol/mg). Our estimate was very close to a previously reported concentration in mouse skeletal muscle (11 pmol/mg)^6^ and of similar magnitude as in different skeletal muscle tissues in rat (25-36 nmol/g) ^7,9,10^. We performed a systematic review of literature but did not find reference values for UDP-GlcNAc for the other tissues.

**Table 1.** UDP-GlcNAc concentration in mouse tissues, pmol/mg tissue

| Tissue | Mean | SD | Sample input<br>(μg initial tissue mass) | Recovery of 0.5 pmol<br>exogenous UDP-GlcNAc (%) |
| --- | --- | --- | --- | --- |
| Liver | 241 | 57 | 3.4 | 109 |
| Kidney | 145 | 37 | 4.6 | 104 |
| Heart | 41.6 | 9.3 | 9.5 | 117 |
| Skeletal muscle<br>(Quadriceps) | 13.7 | 0.8 | 63.5 | 102 |
| Brain<br>(Cerebrum) | 62.6 | 7.4 | 9.9 | 135 |

Brain, n=3; All other tissues, n=4
The exogenous UDP-GlcNAc was spiked into a replicate of the final extract.

### UDP-GlcNAc levels in cultured mammalian cells

Table 2 lists cellular UDP-GlcNAc content in 7 different cell lines: 293T, NIH/3T3, HCT116, AML12, Hepa1-6, HeLa cells, and primary mouse fibroblasts. The UDP-GlcNAc concentrations ranged from 60 to 520 pmol/million cells with the highest per cell content in HeLa cells followed by the liver-originating cells.

**Table 2.**
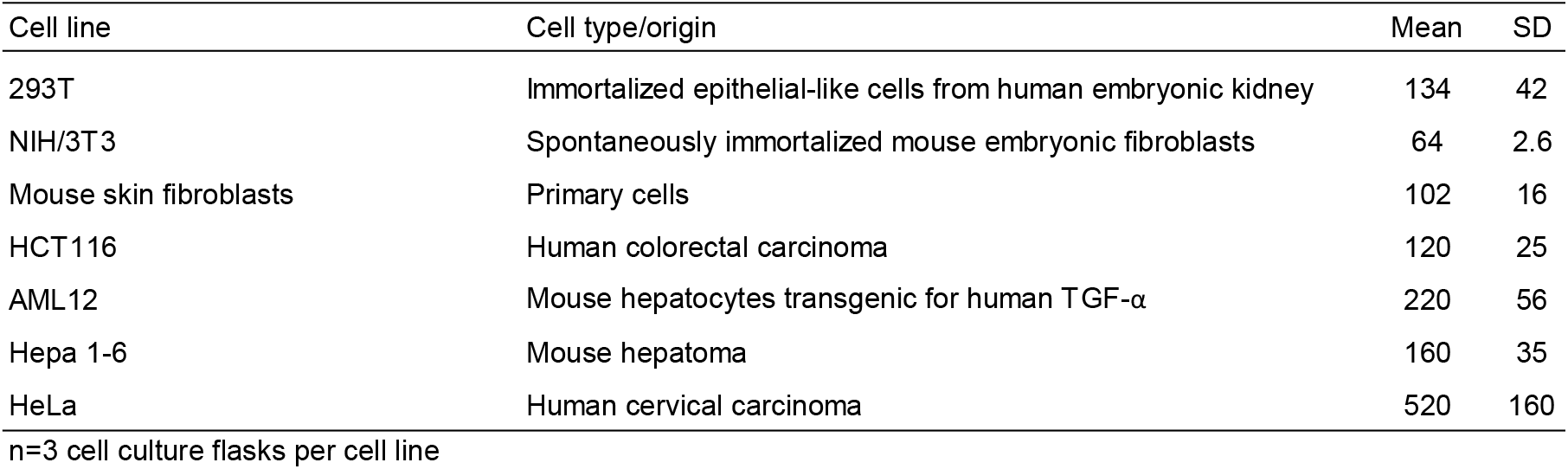
UDP-GlcNAc concentration in cultured mammalian cells, pmol / 10^6^ cells

The HBP and protein O-GlcNAcylation have been targeted by various indirect and direct approaches in published studies, but data on the effect of these manipulations on UDP-GlcNAc levels remain surprisingly scarce. Therefore, we assessed UDP-GlcNAc levels in AML12 hepatocyte cell line subjected to different culturing conditions and metabolic stressors (Figure 3A). Cells grown in 25 or 5 mM glucose had similar UDP-GlcNAc levels, whereas the complete withdrawal of glucose for 16h decreased the UDP-GlcNAc levels by 65%. This depletion was partially prevented by N-acetylglucosamine (GlcNAc) but not by galactose supplementation. In the presence of glucose, GlcNAc elevated the UDP-GlcNAc levels above that of standard culture conditions. Serum-free medium increases hepatocyte-specific gene expression without blocking proliferation in AML12 cells.^26^ Upon serum deprivation, AML12 cells increased their UDP-GlcNAc content (Figure 3A,B).

**Figure 3.**
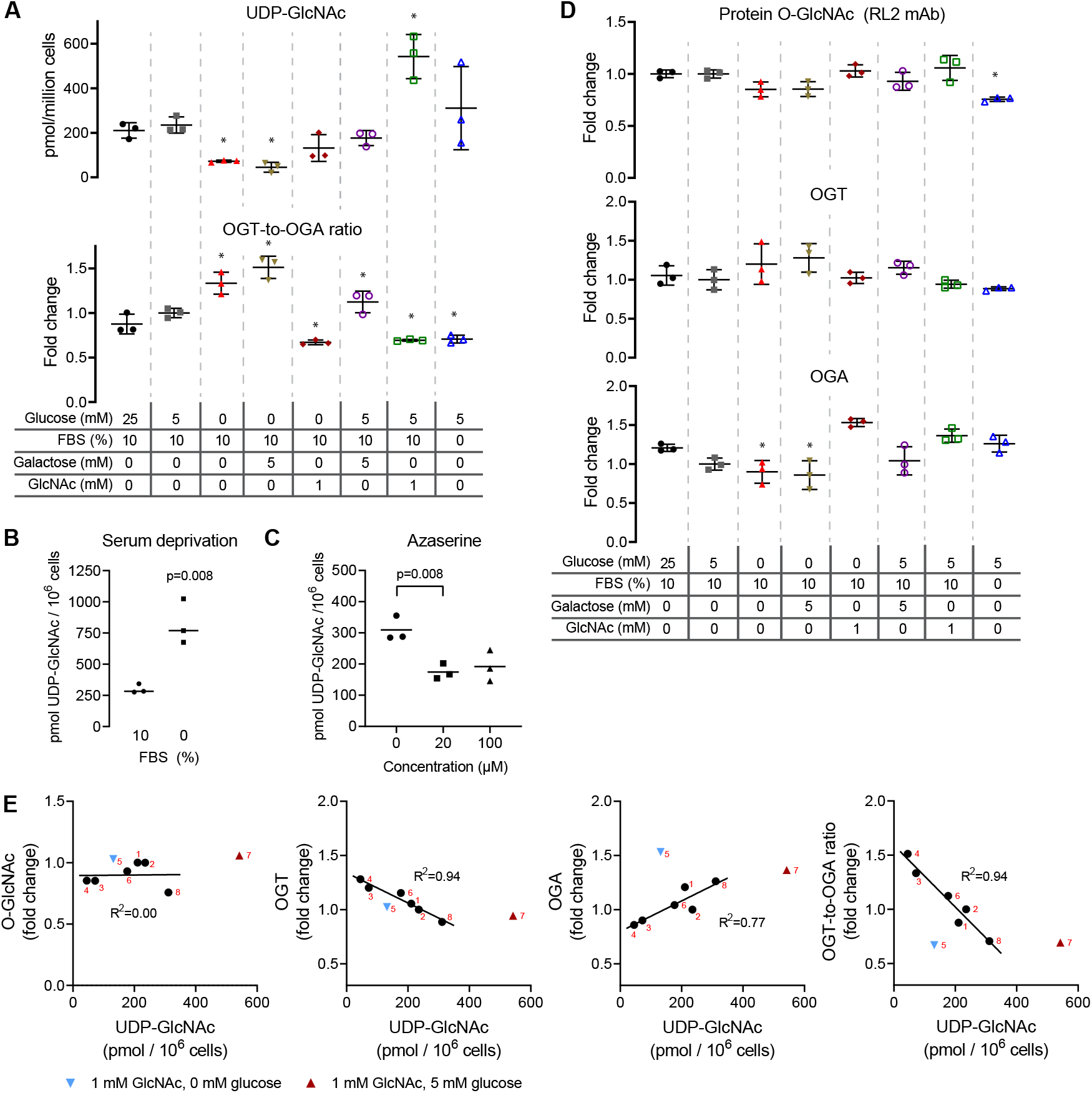
Relationships between UDP-GlcNAc content, protein O-GlcNAcylation, and the expression of OGT and OGA in AML12 cells. (**A**) UDP-GlcNAc concentration and OGT-to-OGA ratio (Western blot) in cells subjected to the indicated culturing conditions for approximately 16h. (**B**) Effect of serum deprivation on UDP-GlcNAc levels. (**C**) Effect of GFAT inhibition (azaserine) on UDP-GlcNAc levels. (**D**) Western blot quantification of protein O-GlcNAcylation, and the expression of OGT and OGA. The samples are same as in figure A. (**E**) Correlations between UDP-GlcNAc concentration and amount of O-GlcNAcylated proteins, and the expression of OGT and OGA, and the OGT-to-OGA ratio. The data points represent average of three replicate cell culture dishes from data in Figure A and D. The small red numerals refer to the experimental groups in Figure A and D with the order from left to right. Data from GlcNAc-supplemented cells (blue and red triangles) were not included in the regression analysis. Supplementary Figure 4 shows representative Western blot detections. Statistics: *, p<0.05 (1-way ANOVA followed Dunnett’s test, 25 mM glucose as control). B and C, unpaired two-sided t-test. The error bars represent ±SD.

Azaserine is a widely used, albeit rather unspecific, inhibitor of the HBP. It inhibits glutamine-dependent enzymes including glutamine fructose-6-phosphate amidotransferases (GFPTs, previously named GFATs) the rate-limiting enzymes of the HBP. In AML12 cells, azaserine decreased UDP-GlcNAc by 50% (Figure 3C), which is similar to that observed in two endothelial cell lines.^27,28^

### Parallel quantification of UDP-GlcNAc, O-GlcNAcylation, and expression of OGT and OGA

Our metabolite extraction procedure allows parallel collection of the total protein fraction. We took advantage of this possibility to measure UDP-GlcNAc and protein O-GlcNAcylation from the same samples. Despite the clear difference in UDP-GlcNAc concentration, the acute (~16h) changes in protein O-GlcNAcylation in the manipulated AML12 cells were minimal (Figure 3A,D,E and Supplementary Figure 4). Cells have been reported to compensate protein O-GlcNAcylation disturbances by rapidly regulating the expression of OGT and OGA.^29,30^ Here, we analyzed the expression patterns of these two enzymes in relation to the UDP-GlcNAc concentration in AML12 cells (Figure 3A,D,E, Supplementary Figure 4). The OGT expression negatively correlated with the cellular UDP-GlcNAc, while the reverse was true for OGA (Figure 3E). The changes were relatively small but given the opposite direction of change, the OGT-to-OGA ratio correlated strongly with the cellular UDP-GlcNAc content. An exception to this correlation came from cells exposed to 1 mM GlcNAc.

In order to examine supraphysiological UDP-GlcNAc levels without GlcNAc supplementation, we generated AML12 cell lines constitutively overexpressing either wild-type (WT) or E328K gain-of-function mutant^31^ GFPT1, an enzyme catalyzing the rate-limiting step in the HBP. Overexpression of the WT enzyme increased UDP-GlcNAc levels approximately 5-fold and the E328K mutant 10-fold (Figure 4A). These supraphysiological UDP-GlcNAc levels led to a relatively meager 1.7-fold increase in O-GlcNAcylated proteins (Figure 4B,D). OGA and OGT expression did not respond to the supraphysiological UDP-GlcNAc availability (Figure 4C,D), suggesting together with the data in Figure 3 that the altered expression of OGA and OGT balance O-GlcNAcylation due to diminished but not elevated UDP-GlcNAc levels in this cell line.

**Figure 4.**
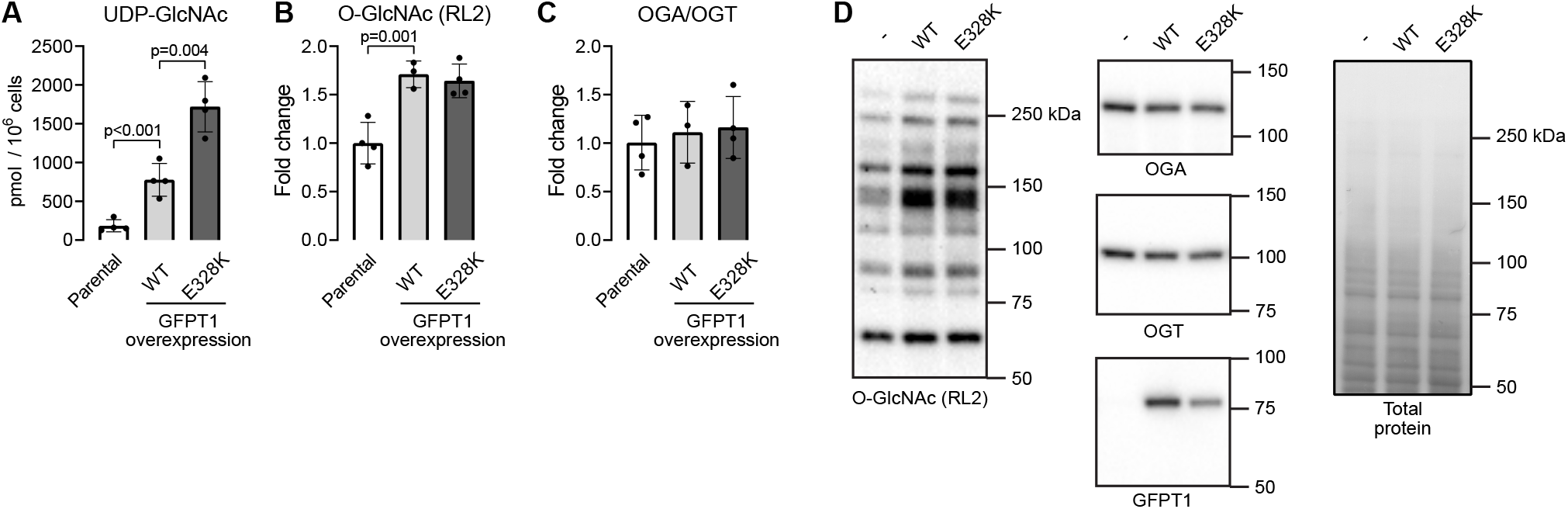
Effect of hexosamine biosynthetic pathway hyperactivity on cellular UDP-GlcNAc and protein O-GlcNAcylation in AML12 cells. (**A**-**C**) UDP-GlcNAc content (**A**), protein O-GlcNAcylation (**B**), and OGA-to-OGT expression ratio (**C**) in parental cells and cells with stable overexpression of wild-type (WT) or E328K mutant GFPT1. (**D**) Representative Western blots and total protein staining as a loading control. Statistics: 1-way ANOVA followed by the selected pairwise comparisons (t-test). The data points represent replicate cell culture flasks. The error bars represent mean and ±SD.

Finally, we measured the same parameters as above from a pancreatic adenocarcinoma cell line (TU8988T) carrying GFPT1 knockout alleles. These cells rely on GlcNAc salvage pathway for generation of UPD-GlcNAc and survive only in the presence of exogenous source of glucosamine.^32^ The parental and the knockout cells had similar very high UDP-GlcNAc content when grown in the presence of 10 mM GlcNAc (Figure 5A, note the logarithmic scale). After 24h of GlcNAc withdrawal, the knockout cells had lost more than 90% of their UDP-GlcNAc. Decrease in O-GlcNAcylated proteins and OGA-to-OGT ratio paralleled the UDP-GlcNAc decline (Figure 5A-D). After 48h, the GlcNAc-starved knockout cells had only ~1% of the baseline UDP-GlcNAc concentration and no detectable O-GlcNAcylated proteins, and notably decreased OGA-to-OGT ratio. On third day of GlcNAc withdrawal, the OGA-to-OGT ratio had further decreased.

**Figure 5.**
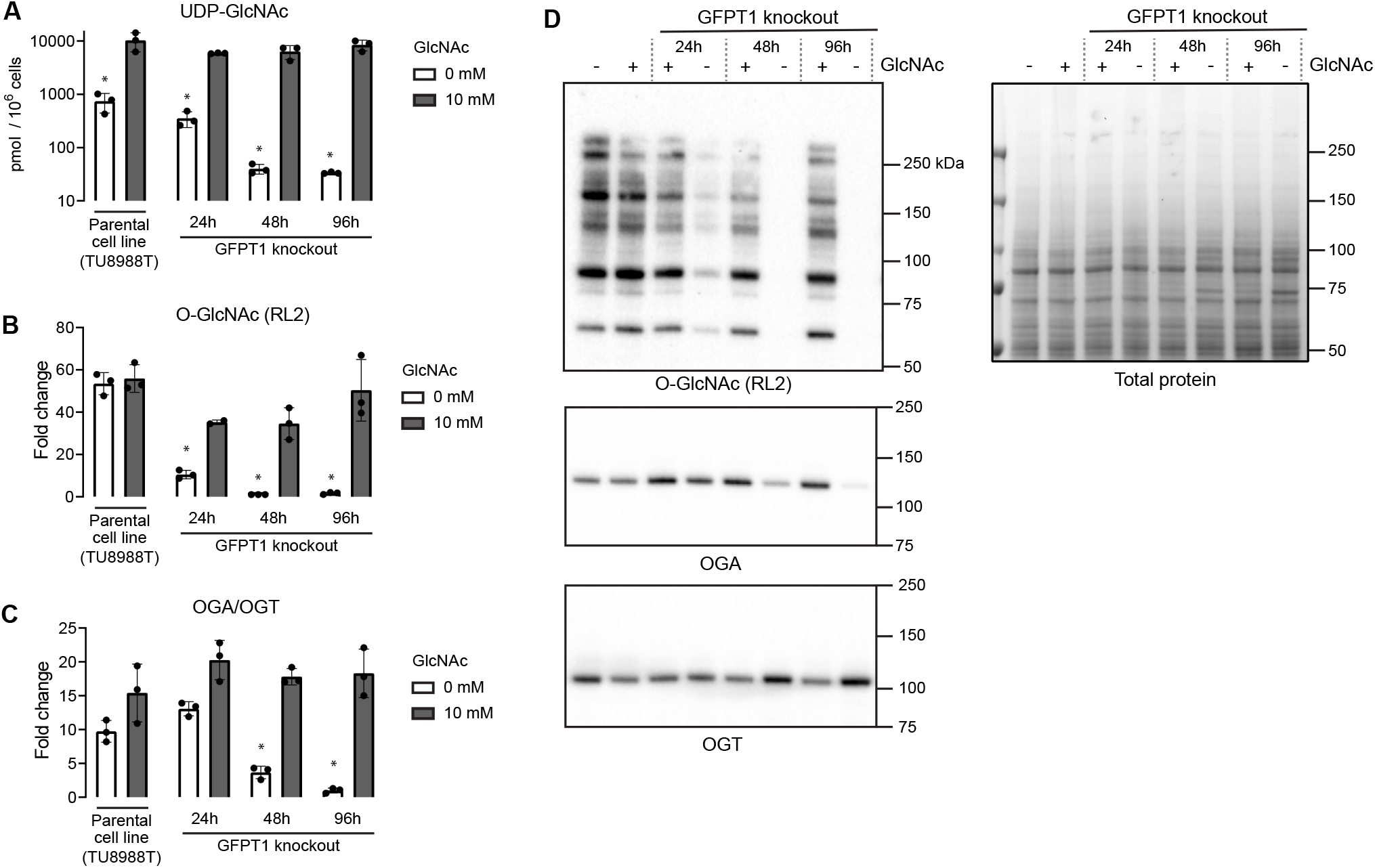
Effect of disrupted hexosamine biosynthetic pathway on cellular UDP-GlcNAc and protein O-GlcNAcylation in a pancreatic adenocarcinoma cell line (TU8988T). (**A**-**C**) UDP-GlcNAc content (**A**), protein O-GlcNAcylation (**B**), and OGA-to-OGT expression ratio (**C**) in parental and GFPT1 knockout cells with and without 10 mM GlcNAc in media. Note the logarithmic y-axis in figure A. The indicated time points refer to duration since replacement of the culture media (start of the GlcNAc starvation). (**D**) Representative Western blots and total protein staining. Statistics: *, Bonferroni-corrected p < 0.0001 (1-way ANOVA followed by the selected pairwise comparisons). The data points represent replicate cell culture flasks. The error bars represent ±SD.

## Discussion

Knowledge on alterations in cellular UDP-GlcNAc concentration is crucial for understanding the mechanisms of O-GlcNAcylation disturbances in different experimental and disease settings. More than 2000 original articles dealing with protein O-GlcNAcylation or the HBP are listed on PubMed, yet we were able to find only a handful of reports showing data on UDP-GlcNAc concentrations. This disproportionate lack of UDP-GlcNAc data is at least partly because, thus far, a simple, practical, and sensitive method to measure this metabolite has not existed. We herein established a robust technique to measure UDP-GlcNAc in any laboratory harboring a typical plate reader and common biochemistry and molecular biology tools. All the reagents are commercially available or, alternatively, the required recombinant enzymes can be produced in-house in bacteria using standard procedures. Moreover, using the sample extraction described here, it is possible to measure UDP-GlcNAc, protein O-GlcNAcylation, and the expression of OGT and OGA from the same samples.

Unexpectedly, we found that global protein O-GlcNAcylation is highly resistant to acute changes in UDP-GlcNAc levels. Two potential reasons for this are the high total cellular concentration of UDP-GlcNAc and the very high affinity of OGT for it.^11–13^ Nevertheless, the expression of OGT and OGA did change upon any decline in UDP-GlcNAc levels, implying rigorous cellular mechanism to sense the UDP-GlcNAc and O-GlcNAcylation statuses. In fact, the OGT-to-OGA ratio proved to be a very sensitive marker of decreased UDP-GlcNAc availability. In contrast, cellular excess of UDP-GlcNAc had less robust effect on the expression of OGT and OGA. Experiments performed with purified proteins suggest that the affinity of OGT for UDP-GlcNAc varies depending on the GlcNAc-acceptor protein.^12^ Thus, the sensitivity of O-GlcNAcylation status of different proteins to disturbances in UDP-GlcNAc concentration may differ. We expect that our method will facilitate tracking down how alterations in cellular UDP-GlcNAc levels affect global and protein-specific O-GlcNAcylation in cell lines and *in vivo*. We also expect that the possibility to quantify UDP-GlcNAc in small biological samples easily will help unravel the underlying causes for altered O-GlcNAcylation in different experimental and disease settings.

The assay described here is highly sensitive with LLOQ of 110 fmol, meaning e.g. that 1 mg tissue biopsy will yield more than enough extract for the assay. There are, however, some limitations. While the dynamic range with purified UDP-GlcNAc standards is wide, we have found that inappropriately diluted biological samples can interfere with the assay. Thus, appropriate sample dilution is important for the absolute quantification of UDP-GlcNAc. Performing the assay takes a full workday. However, although not systematically compared here, all incubation steps can be left to proceed overnight for convenience with relatively minimal effect on the assay performance. For those interested in further development of the assay, the provided Supplementary Table 1 lists some qualitative observations and rationale for the selected assay parameter choices.

In conclusion, we developed a sensitive microplate assay for UDP-GlcNAc, the end-product of the HBP and substrate for protein O-GlcNAcylation. While a large repertoire of tools to probe O-GlcNAcylation have emerged, assessing UDP-GlcNAc has remained difficult for non-specialized laboratories.^33^ Our novel assay fills this methodological gap.

## Methods

### Recombinant proteins

For initial assay development using the dot blotting format, we used commercial recombinant human OGT fragment (Cys323-Glu1041) (R&D systems, 8446-GT). Thereafter, we switched to in-house produced enzyme. A fragment of human OGT cDNA encoding amino acids 322-1041 was PCR-amplified from a human cDNA library (primers MunI-OGT 5’-ATCAATTGACCATGCTGTGTCCCACCCATGCAGA-3’ and OGT-STP-SalI 5’-ATATGTCGACTTATTCAACAGGCTTAATCATGTGGT-3’) and ligated into the EcoRI and XhoI sites of the bacterial expression vector pHAT2, in frame with an N-terminal hexahistidine tag. The plasmid was transformed into the BL21 *E. coli* strain. A single colony from the agar plate was inoculated into 5 ml LB medium containing 100 μg/mL ampicillin and cultured overnight. The bacteria were diluted to 100 ml LB and grown for 3-4 h at room temperature before addition of 0.5 mM IPTG to induce the OGT expression during overnight culture at room temperature. We tested two OGT purification protocols employing nickel or cobalt resin to capture the His-tagged protein. The latter approach yielded slightly purer product (Supplementary Figure 5A) and higher enzymatic activity. This method is described here. Supplementary Materials and Methods describes the alternative purification method. The bacteria were pelleted and suspended into 9 ml lysis buffer comprising 0.5 mg/ml lysozyme, 25 U/ml benzonase, 1 mM MgCl_2_, 0.1% Triton X-100, 0.1 mM TCEP, and 0.5xPBS (5 mM Na-phosphate, 75 mM NaCl, pH 7.4). The bacterial cell walls were digested for 15 min at room temperature and 15 min on ice. Thereafter, the purification was continued while maintaining cold temperature (0-8°C) of the sample and buffers. The lysis was completed by three rounds of 10s probe sonication with 10s cooling time (amplitude 15%, Branson Digital Sonifier 250). Remaining insoluble material was removed by centrifugation (5 min, 20 000g at 4 °C). The supernatant was supplemented to contain 10 mM imidazole, and the phosphate and NaCl concentrations were increased to 20 mM and 300 mM, respectively. This solution was incubated for 1h at ~7°C with 0.25 ml of HisPur Cobalt resin (Thermo Scientific). The beads were allowed to settle to the bottom of a protein purification column and washed 5 times with 10 ml of washing buffer comprising 10 mM imidazole and 0.1% Triton X-100 in 2xPBS. The elution was performed with 150 mM imidazole in washing buffer supplemented with 0.1 mM TCEP. Five 0.25 ml elutes were collected and analyzed by SDS-PAGE (Supplementary Figure 5). The elution fractions 2 to 5 were combined and dialyzed twice (~16h and ~3h at ~7°C) against 200 ml of 50 mM Bis-Tris (Cl^+^) pH 7, 20% glycerol, and 0.1 mM TCEP. The yield was estimated by Bradford assay and BSA standards. The purified OGT was stored in 50% glycerol at −20°C.

The following recombinant proteins were purchased from commercial sources: alkaline phosphatase (Thermo Scientific #EF0651), Antarctic phosphatase (New England BioLabs, #M0289), and human casein kinase II (New England BioLabs, #P6010).

### Preparation of GlcNAc-acceptor peptide-BSA complex

A peptide (NH_2_-KKKY<u>PGGSTPVSSANMM</u>-COOH) containing an O-GlcNAcylation site of human Casein kinase II subunit alpha (underlined sequence) was custom synthesized by Nordic BioSite. A mixture of 6 mM glutaraldehyde, 1 mg/ml peptide, and 1 mg/ml fatty-acid free BSA was incubated in phosphate-buffered saline (PBS) for 30 min to crosslink the peptide to BSA. The reaction was quenched by addition of 12 mM glycine. In our final protocol, the product was sonicated, clarified by centrifugation (3 min 22000g), and centrifuged through a 40 μm cell strainer to increase the uniformity of the coating and to decrease well-to-well variation. The solutions of peptide-BSA complex were stored at −20°C. For some initial experiments, we crosslinked the peptide using 20 mM glutaraldehyde (under the conditions above), quenched the reaction with 19 volumes of 10 mM Tris-HCl pH 7 and removed the uncrosslinked peptide with a Amicon Ultra-4 Centrifugal 10 kDa cut-off column. This approach, however, led to a partial precipitation of the product during centrifugation, and therefore we continued to use the former approach.

### Generation of GFPT1 overexpressing cell lines

The *Mus musculus Gfpt1* cDNA was PCR-amplified (primers: EcoRI-Gfpt1 5’-ATGAATTCGTGACCAACATCATGTGCGG-3’ and Gfpt1-STP-XhoI 5’-ATCTCGAGTTACTCTACTGTTACAGATTTGGC-3’) and the PCR product ligated into pBluescript vector. Internal EcoRI site was removed by site-directed mutagenesis (primers: 5’-CTCATTATTTTTATCAGAGCGCTGG-3’ and 5’-TTCATTGTTATTCATAATGGAATCATCA-3’). The point mutation E328K were introduced by mutagenesis using the following primers: 5’-TCTTCTGCATAAATGAACTAAAGTTG-3’ and 5’-AAATTTTTGAGCAGCCAGAATCTG-3’). After sequencing, the wild type (WT) and E328K variants were subcloned into the Sleeping Beauty transposon donor plasmid ITR\CAG-MCS-IRES-Puro2A-Thy1.1\ITR.

To produce stable cell line overexpressing GFPT1, AML12 cells were co-transfected with the Sleeping Beauty transposon donor plasmid carrying the WT or mutant *Gfpt1* and a plasmid encoding SB100X transposase (1:10 ratio) using FuGENE^®^ HD transfection reagent. Subsequently, the cells were subjected to puromycin selection (2ug/ml). Resistant colonies were pooled and amplified. The expression of GFPT1 was confirmed by Western blot analyses using rabbit anti-GFPT1 antibody (Abcam, EPR4854).

### GFPT1 knockout cells

The TU8988T GFPT1 knockout^32^ and the parental cell lines were a kind gift from Prof. Costas A. Lyssiotis laboratory (University of Michigan, USA).

### Cell culture

AML12 cells were maintained in Dulbecco’s modified Eagle’s medium mixture F12 (DMEM-F12) supplemented with 10% fetal bovine serum, penicillin and streptomycin, 2 mM L-alanyl-L-glutamine, 10 μg/ ml insulin, 5.5 μg/ml transferrin, and 5 ng/ml selenium. Other cell lines were maintained in Dulbecco’s modified Eagle’s medium (DMEM) with 10% fetal bovine serum, penicillin and streptomycin, and 2 mM L-alanyl-L-glutamine. This media was supplemented with 10 mM GlcNAc to maintain the TU89888T GFPT1-knockout cells. All cells were cultured at 37 °C in atmosphere of 5% CO_2_ and 95% air.

For experiments aiming to manipulate UDP-GlcNAc levels, the cells were grown in T-25 cell culture flasks to ~60% confluence using conventional culture conditions. After this, the medium composition was altered as stated in the figures and the figure legends. The cells were washed once with PBS and detached with trypsin and EDTA (TrypLE, ThermoFisher Scientific) and counted. The cells were pelleted by centrifugation, resuspended in 60% MeOH, and stored at −80°C until the extraction.

### Mouse tissue samples

Laboratory animal center of the University of Helsinki maintained the wild-type mice of congenic C57BL/6JCrl background under the internal license of the research group (KEK22-018). Mice of 1 month of age were euthanized by cervical dislocation, tissues immediately excised and placed in liquid N_2_, and stored at −80°C.

For comparison of UDP-HexNAc and UDP-GlcNAc levels, we utilized 23 samples from our other studies comprising liver extracts from mice of four different genotypes: wild-type, *Bcs1l^p.S78G^* heterozygotes and homozygotes, and *Bcs1l^p.S78G^* homozygotes with transgenic expression *of Ciona intestinalis* alternative oxidase.^34^ The genotype differences were irrelevant for this study, and the sole purpose for including these different genotypes was to maximize the number of data points and sufficient variation to allow a meaningful correlation analysis. The transgenic mice were bred under the permit ESAVI/16278/2020 by the State Provincial Office of Southern Finland.

### Preparation of tissue and cell extracts

Frozen pelleted cells or tissue pieces were disrupted in ice-cold 60% MeOH (typically 500 μl per cell pellet or 10 mg tissue) using microtube pestles or roughened glass-to-glass potter homogenizer (for heart, skeletal muscle, and kidney) followed by sonication to complete the homogenization. Further precipitation of proteins was achieved by addition of 225 μl chloroform per 500 μl homogenate and centrifugation (6 min 18000g at 4°C). The aqueous phase was collected and washed three times with ~1.4 ml diethyl ether to remove MeOH. Any remaining layer of diethyl was evaporated by gentle ~10s flow of N_2_ gas. The remaining traces of diethyl ether were evaporated at 65°C in centrifugal vacuum evaporator (SpeedVac Plus SC110A, Savant Instruments) for 5-6 min. The extract volume was estimated by weighing the extract and assuming density of 1 mg/μl. In some initial experiments, chloroform was used instead of diethyl ether to remove MeOH. For data from the TU8988T cells the aqueous-methanol extracts after addition of chloroform were directly evaporated to dryness in miVac centrifugal evaporator (Genevac) without applied heat.

### SDS-PAGE and Western blot

Precipitated proteins from the UDP-GlcNAc extraction were sedimented by addition of 800 μl MeOH and centrifugation (6 min 18000g at 4°C). The protein pellets were dissolved in Laemmli buffer (2% SDS, 4% β-mercaptoethanol, 60 mM Tris-Cl pH 6.8, and 12% glycerol) with the help of sonication and 5 min incubation at 95°C. A modified Bradford reagent containing 2.5 mg/ml α-cyclodextrin to chelate the interfering SDS was employed to measure protein concentrations.^35^ The samples were run on Bio-Rad Criterion TGX gels followed by tank transfer onto PVDF membrane. Equal loading and transfer were verified by Coomassie G-250 staining. After complete destaining of Coomassie G-250, the membrane was blocked with 0.7% Na-caseinate pH 7.4-7.6 and 1% BSA and probed with the following primary antibodies: O-GlcNAc (clone RL2, BioLegend), OGT (ab96718, Abcam), OGA (HPA036141, Merck), and α-tubulin (clone DM1A, Cell Signaling Technology). Peroxidase-conjugated secondary antibodies (#7074 and #7076, Cell Signaling Technology) and enhanced chemiluminescence (ECL) were used for the detection. The ECL reagent^36^ comprised 0.75 mM luminol, 1 mM 4-(Imidazol-1-yl)phenol, 2 mM H_2_O_2_, and 0.05% Tween-20 in 10 mM Tris-Cl buffer pH 9. The luminescence was recorded with Bio-Rad ChemiDoc MP imager.

### UDP-GlcNAc assay in dot blot format

The assay conditions were varied during the method development. Here, the final established assay is described. The O-GlcNAcylation reactions comprised: 10 μg/ml OGT, 25 U/ml alkaline phosphatase, 5 mM Mg-acetate, and 0.2 mg/ml OGT-substrate peptide-BSA complex (concentration based on BSA content), and 50 mM Bis-Tris pH 7.0 (adjusted with HCl). The reactions were carried out in 10 μl volume for 2h at room temperature (21-23°C). The sample volume comprised 2 μl of the reaction mixture. The reactions were stopped by dropping the temperature to 0°C. Bio-Rad microfiltration blotting device was employed to capture the OGT-substrate peptide-BSA complex onto 0.2 μm PVDF membrane (Amersham™ Hybond^®^ P). After completion of the dot blot, the membrane was completely dried, soaked in MeOH, rehydrated, and blocked with 0.7% Na-Caseinate pH 7.4-7.6, 1% BSA for 1h. O-GlcNAcylated residues were detected with the mouse monoclonal RL2 antibody (0.5 μg/ml, 2h), peroxidase-conjugated secondary antibody, and enhanced chemiluminescence.

### UDP-GlcNAc assay in microplate format

The assay parameters were varied and optimized throughout this study. The final established assay conditions are described here. The wells of Nunc^®^ MaxiSorp™ 384 plate were coated with 20 μl of 10 μg/ml OGT-substrate peptide-BSA complex (100 ng BSA) in PBS for ~16h at +4°C. The wells were washed twice with 115 μl of TBS containing 0.05% tween-20 (TBST) and once with 50 mM Bis-Tris buffer pH 7.0. The plate was placed on ice and 16 μl of a reagent mix was quickly added into the wells followed by 4 μl of sample. The final reaction concentrations were 15 μg/ml OGT, 25 U/ml alkaline phosphatase, 5 mM Mg-acetate, 0.3 mg/ml fatty-acid free BSA, and 50 mM Bis-Tris (HCl) pH 7.0. The reactions were caried out for 2-3h at room temperature. The wells were washed twice with TBST and once with TBS. Then, the wells were incubated with 20 μl of RL2 antibody (0.5 μg/ml) for 1-2h. Following 5 washes with TBST and one wash with TBS, 20 μl of peroxidase-conjugated secondary antibody (1:3000, #7076, Cell Signaling Technology,) was added into each well and incubated for 0.5-1h. Both antibodies were diluted in 1% BSA in TBST (0.2% Tween-20). After repetition of washing steps, 12.5 μM Amplex UltraRed (Invitrogen, #A36006) and 1 mM H_2_O_2_ were utilized as the peroxidase substrates in 100 mM potassium phosphate buffer pH 6.7. The chemifluorescence was developed for 45 min in dark and the end-point fluorescence read upon 530 nm excitation and 590 nm emission with BioTek Synergy H1 plate reader. To prevent evaporation during the incubation steps the plates were sealed with an adhesive membrane.

### UPLC-MS-based quantification of UDP-HexNAc

To approximately 15 mg of snap-frozen liver samples, 600 μl of ice-cold chloroform-methanol (2:1) solution was added. The tissue pieces were ground with a ball mill and subjected to three 10-min ultrasonic bath treatments on ice and three freeze-thaw cycles (−196°C to 4°C). Samples were vortexed in cold room (4°) for 30 min prior to addition of 400 μl milli-Q water. After addition of water, samples were further vortexed for 30 min and centrifuged to induce phase separation. Upper water-methanol phase was transferred to a new Eppendorf tube and evaporated to dryness in vacuum concentrator (Genevac, miVac Duo). Samples were reconstituted in 100 μl water and analyzed with UPLC-QTRAP/MS (ABSciex).

Chromatographic separation was performed in Waters Premier BEH C18 AX column (150 x 2.1mm, ø 1.7μm) at 40°C. Elution solvents were 10 mM ammonium acetate in water, pH 9.0 (A) and acetonitrile (ACN) (B) with flow rate of 0.3 ml min^-1^. Linear gradient started with 98% A and decreased to 30% in 3 min, then to 10% in 1 min, back to 98% in 4.01 min time, and stabilized for 1 min, with 5 min total runtime. Injection volume was 1 μl. UDP-HexNAc was analyzed with UPLC-6500 + QTRAP/MS (ABSciex) in negative ion mode. Retention time was 1.32 min. Multiple Reaction Monitoring (MRM) method with two transitions, quantitative and qualitative were performed: quantitative 606 → 385, and qualitative 606 → 159.

After the UPLC-MS runs, the extracts were evaporated to dryness, and stored (~1.5 years) at −80°C, before measurement of UDP-GlcNAc with the enzymatic method.

### Statistical analyses

The signal-to-noise ratio was defined as background-subtracted fluorescence divided by the standard deviation of the background (n>6). LLOQ and LLOD were estimated empirically (variation and standard curve fit) and mathematically based on signal-to-noise cut-off of 10 and 2, respectively. Signal-to-background was defined as signal divided by the background (0 nM UDP-GlcNAc). The bar graphs for data from biological replicates represent mean and standard deviation (SD). Mean and standard error of mean (SEM) are shown for technical replicates. GraphPad Prism (version 9.3.1) was utilized for non-linear curve fitting and other statistical analyses. Differences between the groups were assessed with two-tailed t-test or 1-way ANOVA followed by an appropriate multiple comparisons tests as stated in the figure legends.

### Step-by-step protocol

Detailed protocols are provided as supplementary files.

## Supporting information

Step-by-step protocol - sample extraction

step-by-step protocol UDP-GlcNAc assay

## Acknowledgements

We thank Prof. Costas Lyssiotis and Dr. Lin Lin (University of Michigan, USA) for providing the GFPT1 knockout cell lines and for valuable advice regarding their culture. The *Methanococcus maripaludis* UDP-GlcNAc dehydrogenase bacterial expression construct was a kind gift by Dr. David E. Graham (Oak Ridge National Laboratory, USA). Part of the work was carried out with the support of HiLIFE Laboratory Animal Center Core Facility, University of Helsinki, Finland. We acknowledge the funding from Folkhälsan Research Center (V.F. and J.K.), Jane and Aatos Erkko Foundation (J.K.), the Foundation for Pediatric Research (V.F.), Finska Läkaresällskapet (V.F.), The Liv och Hälsa Foundation (V.F.), Magnus Ehrnrooth Foundation (M.S.), and Alfred Kordelin Foundation (J.P.). The graphical illustrations were created with the help of BioRender.com.

## Author contributions

J.P. invented the assay concept and wrote the first manuscript draft. J.K. was responsible for cloning of the OGT construct. J.K., M.S., J.P. took part in the production of the recombinant OGT. J.P., M.S., and R.B. performed the UDP-GlcNAc measurements. M.S., D.U., and R.B. were responsible for the cell culture work. D.U. cloned the GFPT1 plasmids and generated the GFPT1-overexpressing cell lines. N.S. performed the UPLC-MS measurements. All authors critically read and commented the manuscript, and J.P. and J.K. revised it accordingly. J.P. and J.K. supervised the study.

---

**Supplementary Figure 1.**
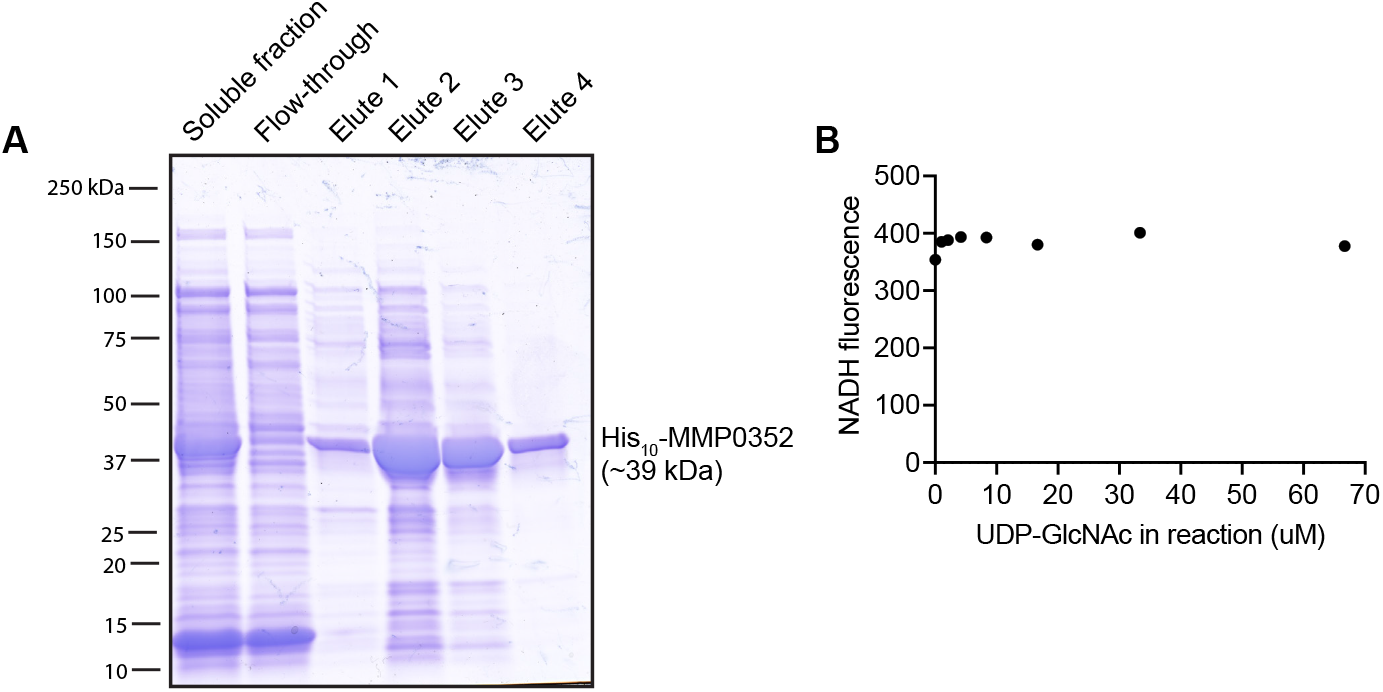
Testing of applicability of *Methanococcus maripaludis* UDP-GlcNAc dehydrogenase (MMP0352) for quantification of UDP-GlcNAc. (**A**) SDS-PAGE of purified recombinant His-tagged UDP-GlcNAc dehydrogenase (elute fractions 1 to 4). (**B**) Testing of varying concentrations of UDP-GlcNAc for NADH generation by the UDP-GlcNAc dehydrogenase. The measurements were performed as described by Namboori and Graham (18). The enzyme was expressed in *E. coli* BL21 (DE3) (pDG441) cells, purified by Ni^2+^ affinity chromatography, and dialyzed to PBS. The bacterial expression construct was a kind gift by Dr. David E. Graham (Oak Ridge National Laboratory, USA).

**Supplementary Figure 2.**
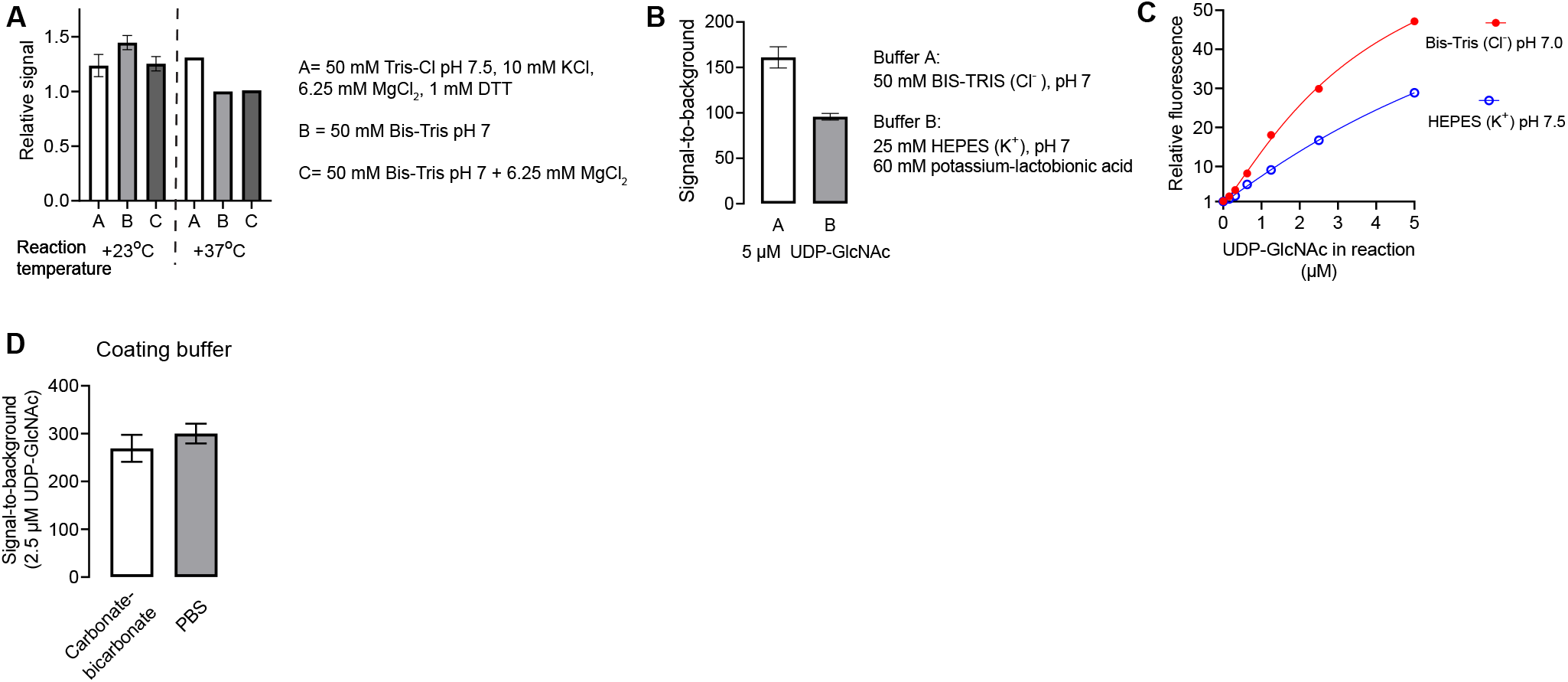
Optimization of the assay. (**A**-**B**) Data produced using the dot blot format of the assay. All reactions contained 5 μM UDP-GlcNAc, 10 μg/ml OGT, and 25 U/ ml alkaline phosphatase. (**A**) Effect of O-GlcNAcylation reaction temperature and buffer on the obtained signal. (**B**) Comparison of the indicated buffers. (**C**-**D**) Optimization data produced using the microplate format. (**C**) Comparison of Bis-Tris and HEPES-based assay buffers. (**D**) Effect of coating buffer. The bar graphs represent mean and SEM of technical triplicates.

**Supplementary Figure 3.**
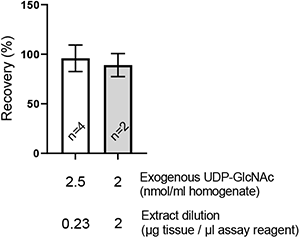
Recovery of exogenous UDP-GlcNAc added to mouse kidney tissue homogenates. In this experiment, the residual MeOH in the extract was removed by diethyl ether washes followed by the brief vacuum evaporation of the organic solvent traces. The sample dilution refers to the original tissue mass worth of polar metabolite extract in the assay. The bars graphs represent mean and SEM.

**Supplementary Figure 4.**
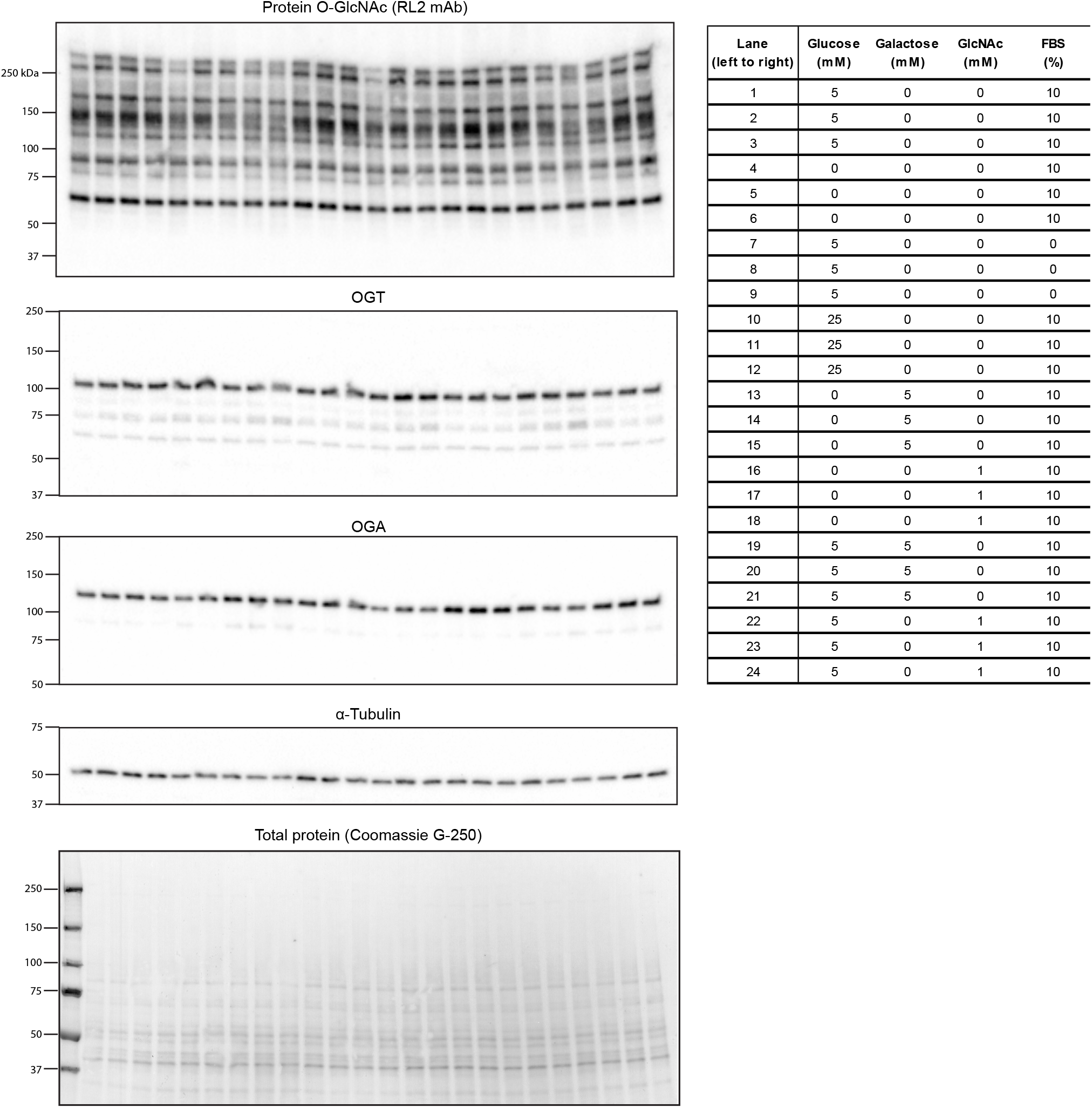
Western blots used for quantifications in Figure 4. α-tubulin and on-membrane total protein staining are shown as loading controls.

**Supplementary Figure 5.**
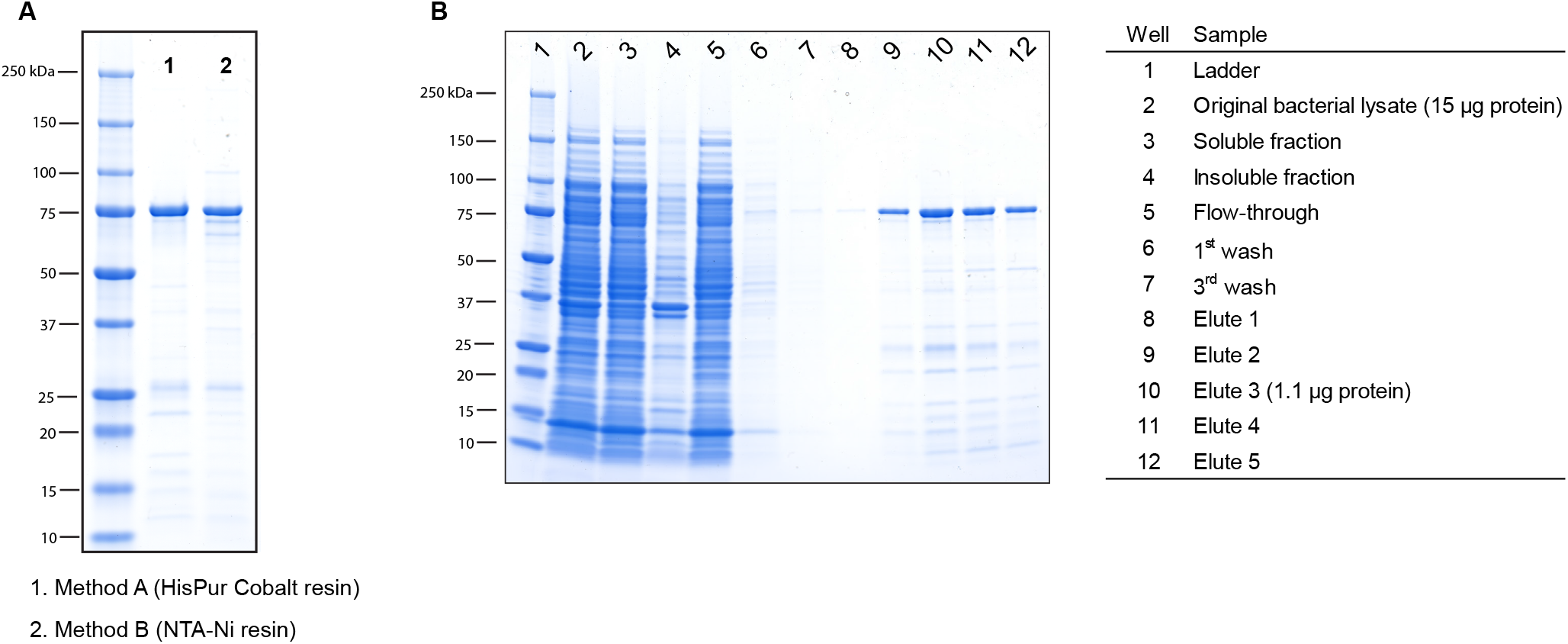
SDS-PAGE analysis of purified recombinant OGT. (**A**) Comparison of OGT purified using cobalt resin (Materials and Methods) and NTA-Ni resin (Supplementary Materials and Methods). (**B**) Representative SDS-PAGE analysis of fractions from Cobalt resin purification of the His-tagged OGT.

**Supplementary Table 1.**
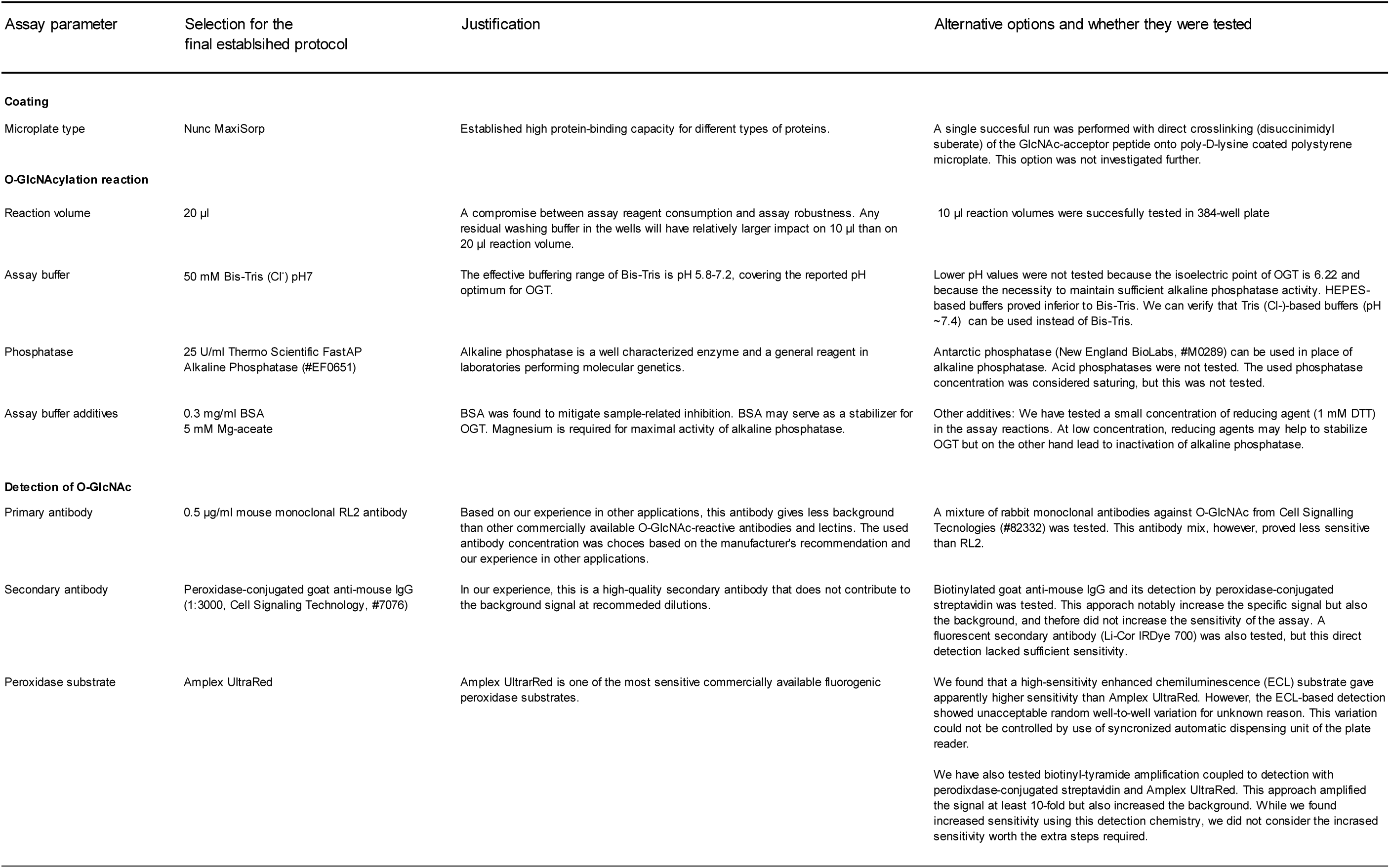
Qualitative observations and rationale for the selected assay parameters

