## Supplementary material for "Enzymatic assay for UDP-GlcNAc and its application in the parallel assessment of substrate availability and protein O-GlcNAcylation": Step-by-step protocol - sample extraction

Supplementary material, Sunden et al. 2023

### Step-by-step protocol:
### Extraction of polar metabolites and total protein fraction

#### Reagents

- Ice-cold 60% MeOH:

- Chloroform. Volatile and toxic, use fume hood! Do not use serological pipettes made of polystyrene to dispense chloroform!

- Diethyl ether. Volatile and toxic, use fume hood! Do not use serological pipettes made of polystyrene to dispense diethyl ether!

- Milli-Q-grade water

- 1.5 ml microtube

- 2 ml microtubes

- Microtube homogenizer for soft tissues (e.g. Nippon Genetics cat. no. NG010).

- Tissue grinder for fibrous tissues (e.g. WHEATON® Tenbroeck Tissue Grinder, WVR cat. no. 62400-493).

- Microtip probe sonicator.

- SDS buffer: 2% SDS, 4% β-mercaptoethanol, 12% glycerol, and 60 mM Tris-Cl pH 6.8.

#### Extraction of polar metabolites from snap-frozen tissue

1. Cut ~10 mg of frozen tissue into a 1.5 ml microtube. Keep the samples dry ice cold.

2. Homogenize soft tissues (e.g. liver and kidney) in 0.5 ml of ice-cold 60% MeOH using a microtube pestle homogenizer. Place the homogenized sample back on dry ice before proceeding to the next sample.

   For fibrous tissues (e.g. skeletal muscle), we recommend a more harsh homogenization method e.g. roughened glass-to-glass potter homogenizer.

3. Sonicate the homogenate for 12s with a probe sonicator and appropriate amplitude setting for the volume. Place the sonicated sample back on dry ice or continue immediately to step 4.

4. Place the samples on wet ice. Add 225 µl chloroform.

5. Vortex 10s. Shake vigorously for 30s.

6. Centrifuge 3 min 18000g at 0-4°C.

7. [Optional] Discard most of the lower phase (chloroform-MeOH) with a syringe (same syringe can be used for all samples). Centrifuge 3 min 18000g at 0-4°C. This step allows almost complete collection of the upper phase (MeOH-$H_2O$) in the next step.

8. Collect the upper MeOH-$H_2O$ phase into a new tube.
   Use pre-weighted 2 ml microtubes if continuing with option 2 in the next step.

   [Optional] Save the interphase and the remaining lower phase for collection of proteins.

Option 1

Evaporate the extracts to dryness in a centrifugal vacuum evaporator that can dry the extracts without applied heat within a reasonable time (~1h).

Option 2

Wash the aqueous fractions with diethyl ether to remove residual MeOH.

a) Add 1.4 ml of diethyl ether.
   Vortex for ~5s.
   Briefly centrifuge (e.g. 1s 14000g).
   Discard most of the upper layer.

b) Repeat step a) three times. Approximately 200 µl of aqueous extract should be remaining.

c) Evaporate any residual diethyl ether layer under $N_2$ gas flow for a few seconds. If a source of $N_2$ gas is not available, keep the sample tubes gaps open at room temperature until the remaining diethyl ether layer has evaporated (a few minutes).

d) Evaporate the remaining traces of diethyl ether in a centrifugal vacuum evaporator for 5 min. If a centrifugal vacuum evaporator is not available, place the sample tubes, gaps open, into a heat block preheated to 90°C for 3 min.

e) Weigh the remaining amount of liquid (1 mg ~ 1 µl) to estimate the volume of the extract.

Option 3

Due to the high sensitivity of the final optimized assay, it is very likely that in the case of many sample types, the extracts from step 8 can be directly diluted beyond any interference by MeOH.

9. Store the extracts at -80°C.

#### Extraction of polar metabolites from cultured cells

##### Option 1

a. Detach, count, and pellet the cells using standard procedures.
b. Resuspend pelleted cells in 0.5 ml ice-cold 60% MeOH.
c. Continue from step 3 of the extraction protocol for tissue samples.

The option 1 allows normalization of UDP-GlcNAc concentrations to either cell number of protein amount.

##### Option 2

a. Wash the cells once with ice-cold PBS.
b. Scrape the cells directly in an appropriate volume of ice-cold 60% MeOH (0.5 ml per 6 cm dish)
c. Continue from step 3 of the extraction protocol for tissue samples. Adjust the chloroform volume based on the volume of 60% MeOH.

We recommend the option 2 especially if more labile metabolites than UDP-GlcNAc (e.g. nucleotides) are to be measured from the same extract. With option 2 the UDP-GlcNAc concentrations can be normalized to protein amount.

#### Collection of total protein fraction

1. Add 800 µl of ice-cold 100% MeOH to the chloroform-interphase fraction from step 8.
   Mix.

2. Centrifuge 3 min 18000g at 0-4°C
   Remove the supernatant.

3. Centrifuge 1s 18000g.
   Remove any remaining liquid.

4. Add an appropriate volume of SDS buffer (typically 400 µl for 10 mg tissue samples or 150 µl for 6 cm cell culture dishes).

5. Let the protein pellets stand in SDS buffer for 10 – 30 min at room temperature.

6. Sonicate to completely dissolve the proteins.

7. (Optional) Incubate 5 min at 95°C.

8. Measure protein concentration with a method that is compatible with the SDS buffer.
   We recommend a modified SDS-compatible Bradford assay (ref. 35 in the Reference list).

9. The protein fractions are ready for Western blot analyses after addition bromophenol blue.
