## Supplementary material for "Enzymatic assay for UDP-GlcNAc and its application in the parallel assessment of substrate availability and protein O-GlcNAcylation": step-by-step protocol UDP-GlcNAc assay

Supplementary material, Sunden et al. 2023

### Step-by-step protocol:
### Quantification of UDP-GlcNAc

#### Reagents and materials

- Phosphate-buffered saline (PBS)

- OGT substrate peptide: $NH_2$-KKKYPGGSTPVSSANMM-COOH
  1 mg/ml in PBS store in aliquots at -20°C

- 10 mg/ml fatty acid-free BSA in water. Store at -20°C.
  The exact concentration should be determined by reading 280 nm absorbance and employing the extinction coefficient of 0.667 $mg^{-1}*ml*cm^{-1}$.

- 0.6 M glutaraldehyde in water. Store at -20°C.

- 0.6 M glycine. Store at -20°C.

- Preparation of 2 mg/ml OGT-substrate peptide-BSA complex (1 mg/ml BSA, 1 mg/ml peptide).

1. Mix:
   0.2 ml of 1 mg/ml OGT substrate peptide in PBS
   2 µl of 100 mg/ml BSA
   2 µl of 600 mM glutaraldehyde
2. Incubate 0.5h at room temperature (~22°C).
3. Add 4 µl of 600 mM glycine (f.c. 12 mM).
4. Sonicate two times 10s on ice with at least 10s pause between the sonication steps.
5. Centrifuge 18000-22000g 3 min. Collect the supernatant.
6. Centrifuge the supernatant (800g 1s) through a 40 µm cell strainer.
7. Aliquot and store at -20°C

- MaxiSorp high-protein binding 384-well microplate with black wells.

- UDP-GlcNAc sodium salt (Sigma, U4375)
  Dissolve 25 mg in 384 µl water to obtain a 100 mM stock solution.
  Prepare a working solution of 1 mM. Aliquot and store the solutions at -20°C.

- UDP-GlcNAc standard curve:
  Prepare an 8-point 1:2-serial dilution covering 1 to 0 µM.
  Store at -20°C.

- Recombinant human OGT fragment (0.35 mg/ml, batch dependent).
  Commercially available from R&D systems (cat. no. 8446-GT).
  See Materials and Methods for in-house production of the enzyme.

- 100 mM Bis-Tris pH 7.0 (adjust with HCl)

- 500 mM Mg-acetate

- 1 kU/ml alkaline phosphatase (Thermo Scientific #EF0651)

- Tris-buffered saline (TBS)
- TBS + 0.1% tween-20 (TBST, wash buffer).
- Antibody diluent: 1% BSA in TBS + 0.2% tween-20

- RL2 mouse monoclonal antibody (Biolegend, #100102)

- Peroxidase-conjugated anti-mouse IgG (Cell Signaling Technologies, #7076)

- 100 mM potassium phosphate buffer pH 6.7

- 10 mM Amplex UltraRed (Invitrogen, A36006)
  Dissolve 1 mg in 333 µl of DMSO. Aliquot and store at -20$^{o}$C. Minimize exposure to light.

- Stabilized ~3% hydrogen peroxide solution.
  Make an appropriate dilution (e.g. 1:50) in water and determine the exact concentration of the stock solution by measuring 240 nm absorbance and employing the extinction coefficient of 43.6 $M^{-1}cm^{-1}$.

- Assay reagent.
  Recipe for 100 reactions with the final reaction volume of 20 µl (16 µl reagent and 4 µl sample):
  1000 µl of 100 mM Bis-Tris pH 7.0
  384 µl of $H_2O$
  60 µl of 10 mg/ml fatty-acid free BSA
  20 µl of 500 mM Mg-acetate
  50 µl of of 1 kU/ml Alkaline Phosphatase
  86 µl of 0.35 mg/ml OGT

Final concentrations (after addition of sample): 15 µg/ml OGT, 25 U/ml Alkaline phosphatase, 50 mM BIS-TRIS (HCl) pH 7, 5 mM Mg-acetate, and 0.3 mg/ml BSA

- Amplex Ultra Red HRP substrate.
  Recipe for 10 ml:
  10 ml 100 mM potassium phosphate buffer pH 6.7
  12.5 µl of 10 mM Amplex UltraRed
  $H_2O_2$ to final concentration of 1 mM.

  Prepare directly before use! Minimize light exposure. Do not place this solution in a reagent reservoir that have been exposed to peroxidase-conjugated antibodies.

**Measurement of UDP-GlcNAc**

Do not allow the plate to completely dry at any point. The O-GlcNAcylation reactions should be set up on ice if a multichannel pipette is not used. All antibody solutions and Amplex UltraRed solution should be prepared in buffers equilibrated to room temperature to minimize temperature gradients. Washing steps should be performed similar to in ELISA assays: pouring off most of the buffer and tapping the plate against paper towels.

1. Add 20 µl of 10 µg/ml OGT-substrate peptide-BSA complex in PBS into the wells. Seal the plate and spin down. Incubate overnight at +4$^{o}$C.

2. Wash twice with TBST (115 µl)
3. Wash once with 50 mM Bis-Tris pH 7.

4. Discard the buffer and place the plate on ice.

5. Add 16 µl of ice-cold assay reagent.
6. Add 4 µl of sample
   Seal the plate, mix gently by tapping, and spin down.

7. Incubate 2-3h at room temperature (20-25 $^{o}$C).
   The plate can be placed on top of a metal bock of suitable size to speed up warming of the plate and to minimize temperature gradients.

8. Wash twice with TBST
   Wash once with TBS.
   The last wash with TBS prevents formation of bubbles in the wells.

9. Add 20 µl of RL2 mAb 1:1000 in 1% BSA in TBS + 0.2% tween-20 with a multichannel pipette.
   Seal the plate and spin down.
   Incubate 1-2h at room temperature.

10. Wash 5 times with TBST
    Wash once with TBS

11. Add 20 µl of anti-mouse secondary antibody 1:3000 in 1% BSA in TBS + 0.2% tween-20 with a multichannel pipette.
    Incubate 0.5-1h at room temperature.

12. Wash 5 times with TBST.
    Leave the wells in TBS

13. Prepare Amplex UltraRed HRP substrate.
    Empty the wells.
    Add 20 µl Amplex UltraRed HRP substrate with a multichannel pipette.
    Seal the plate and incubate 45 min in dark.

14. Remove the plate seal. Measure fluorescence upon 530 nm excitation and 590 nm emission.
