## Supplementary Materials and Methods for "Enzymatic assay for UDP-GlcNAc and its application in the parallel assessment of substrate availability and protein O-GlcNAcylation"

### Competitive enzyme-linked lectin-binding approach to measure UDP-GlcNAc

This is a description of tested alternative approach to measure GlcNAc or UDP-GlcNAc. The principle of the assay is that GlcNAc or UDP-GlcNAc competes with GlcNAc moieties of glycoproteins for binding to wheat germ agglutinin (WGA). We did not continue to develop this method further because of the complex assay nature and lack of robustness and sensitivity. Nunc MaxiSorp-96 plate were coated overnight with 5% skimmed milk as a source of glycoproteins. Per each sample to be analysed, a 9-point 1:2 dilution series of biotinylated WGA covering 5 µg/ml to 20 ng/ml were distributed into the wells of the microplate (150 µl volume). The diluent for WGA was TBS containing 1% tween-20, 1% polyvinylpyrrolidone, 1% polyvinyl alcohol, and 0.4% polyethylene glycol (4 kDa), 1 mM $MgCl_2$, and 1 mM $CaCl_2$. GlcNAc standard samples (50 µl volume) were added into each WGA dilution series. The wells were washed 5 times with TBST. The biotinylated WGA remaining in the wells was detected with peroxidase-conjugated streptavidin and Amplex UltraRed.

### Ni-NTA affinity purification of His-tagged OGT.

The induced bacteria were harvested and suspended into lysis buffer comprising 50 mM sodium phosphate pH 8, 300 mM NaCl, 10 mM of imidazole, and 1 mg/mL of lysoszyme. After 20 min incubation at room temperature, the bacterial suspension was sonicated to complete the lysis. The lysate was centrifuged (25 min, 12 000g at 4 ˚C) and the supernatant mixed with 1 ml of 50 % Ni-NTA bead slurry for 60 min at 4 ˚C. The beads were collected into a protein purification column and washed twice with the lysis buffer. The elution was performed with 300 mM imidazole (in 50 mM sodium phosphate, 300 mM NaCl). Four 0.5 ml elutes were collected and analyzed by SDS-PAGE. The elution fractions were combined and supplemented with 0.1% Triton X-100. The buffer was changed to 100 mM Bis-Tris pH 7, 1 mM DTT by dialysis. The yield was estimated by Bradford assay and BSA standards. The purified OGT preparation was aliquoted and snap-frozen in liquid nitrogen and stored at -80ºC or stored as a 50% glycerol stock at -20ºC.
